# *Bacteroidales* species are a reservoir of phase-variable antibiotic resistance genes in the human gut microbiome

**DOI:** 10.1101/2021.04.26.441444

**Authors:** Wei Yan, A. Brantley Hall, Xiangfang Jiang

## Abstract

Phase-variable antibiotic resistance genes (ARGs) can mitigate the fitness cost of maintaining ARGs in the absence of antibiotics and could potentially prolong the persistence of ARGs in bacterial populations. However, the origin, prevalence, and distribution of phase-variable ARGs remains poorly understood. Here, we sought to assess the threat posed by phase-variable ARGs by systematically searching for phase-variable ARGs in the human gut microbiome and examining their origin, prevalence, and distribution. Through metagenomic assembly of 2227 human gut metagenomes and genomic analysis of the Unified Human Gastrointestinal Genome (UHGG) collection, we identified phase-variable ARGs and categorized them into three classes based on the invertase regulating phase variation. In the human gut microbiome, phase-variable ARGs are commonly and exclusively distributed in *Bacteroidales* species. Through genomic analysis, we observed that phase-variable ARGs have convergently originated from ARG insertions into phase-variable capsule polysaccharide biosynthesis (CPS) loci at least three times. Moreover, all identified phase-variable ARGs are located within integrative conjugative elements (ICEs). Therefore, horizontal transfer via ICEs could explain the wide taxonomic distribution of phase-variable ARGs. Overall, these findings reveal that phase-variable CPS loci in *Bacteroidales* species are an important hotspot for the emergence of clinically-relevant phase-variable ARGs.

## INTRODUCTION

The proliferation of antibiotic resistance genes (ARGs) has compromised antibiotic treatment for bacteria infections (1). The human gut microbiome is an important reservoir of ARGs (2–4) and the spread of ARGs from gut microbes to pathogens has been documented (5). Therefore, ARGs in the human gut microbiome pose a growing threat to human health.

Often, bacteria carrying ARGs are outcompeted by susceptible strains due to the costs associated with the maintenance and expression of the ARGs (6–8). Though it is costly, bacteria can ameliorate the fitness costs of maintaining ARGs through different strategies (9), such as no-cost, low-cost or gain of fitness mutations (10, 11), compensatory mutations at a second site (12–14), or genetic co-selection of resistance genes in genetic linkage (15, 16). Phase-variable ARGs, which were only recently reported (17), are a newly-identified mechanism for antibiotic resistant bacteria to mitigate the fitness cost of encoding ARGs.

Phase variation refers to a reversible change that generates phenotypic variation which helps bacteria adapt to rapidly changing environments (18, 19). Phase variation often manifests through reversible inversion of DNA regions containing promoters such that in one orientation a downstream gene is expressed while in the alternate orientation, the downstream gene is not expressed (17). Such DNA inversions are generally mediated by invertases, which recognize inverted repeats flanking the invertible region and catalyze the reversible inversion (20–22). Phase-variable genes often regulate characteristics important for bacterial colonization and virulence including fimbriae (23, 24), flagella (25), and capsular polysaccharides (CPS) (26, 27).

Recent advances in computational methods have contributed to the effective identification of the intergenic invertible DNA regions in microbial genomes (17, 28). ARGs were found to be regulated by invertible promoters in certain human gut bacteria (17). Resistant bacteria with phase-variable ARGs could maintain these ARGs for a longer period of time in the microbial community, by switching the expression of ARGs to the OFF orientation in the absence of antibiotics to ameliorate the fitness cost (17). The emergence of phase-variable ARGs increases the burden for humans to combat antibiotic resistance and the spread of phase-variable ARGs to pathogens poses an increasing threat to human health. Therefore, we need a better understanding of the evolution of phase-variable ARGs as well as their taxonomic and geographic distribution to combat them.

Here, we took advantage of the considerable amount of human gut metagenomic data generated during the last decade to examine the origin, evolution and prevalence of phase-variable ARGs on a large scale. We systematically searched for phase-variable ARGs through metagenomic assembly of 2227 human gut metagenomes and the Unified Human Gastrointestinal Genome (UHGG) collection of human gut genomes. We found that phase-variable ARGs were commonly distributed in *Bacteroidales* species. Genomic analysis showed that phase-variable ARGs have been convergently derived from ARG insertions into phase-variable CPS loci. Notably, the identified phase-variable ARGs were found to have been mobilized through ICEs and have been widely geographically distributed. Our results reveal the prominent role of phase-variable CPS loci in *Bacteroidales* species in providing hotspots for the emergence of clinically-relevant phase-variable ARGs.

## RESULTS

### Identification and classification of phase-variable antibiotic resistance genes

To expand the known repertoire of phase-variable ARGs, we searched for phase-variable ARGs from publically available human gut metagenomic datasets, comprising a total of 2227 samples, using the tool PhaseFinder (Table S1). Briefly, we assembled the metagenomic data then searched for phase-variable regions with PhaseFinder by aligning the unassembled reads back to the metagenomic assemblies. Then, phase-variable ARGs were identified by scanning regions downstream of an identified phase-variable region for ARGs. In total, this search uncovered 61 contigs containing phase-variable ARGs. In all phase-variable ARGs identified, the invertible region was located immediately downstream of a gene encoding an invertase. We identified putative promoter sequences in the invertible region of all 61 phase-variable regions. Therefore, we concluded that these 61 ARGs are likely regulated by invertible promoters.

To analyze the origin and evolution of phase-variable ARGs, we constructed a phylogenetic tree based on the nucleotide sequence of the invertases. We grouped the invertible promoters regulating ARGs (IP-ARGs) into three distinct classes, denoted IP-ARG-1, IP-ARG-2, and IP-ARG-3 (Figure 1, Table S2).

**Figure 1:**
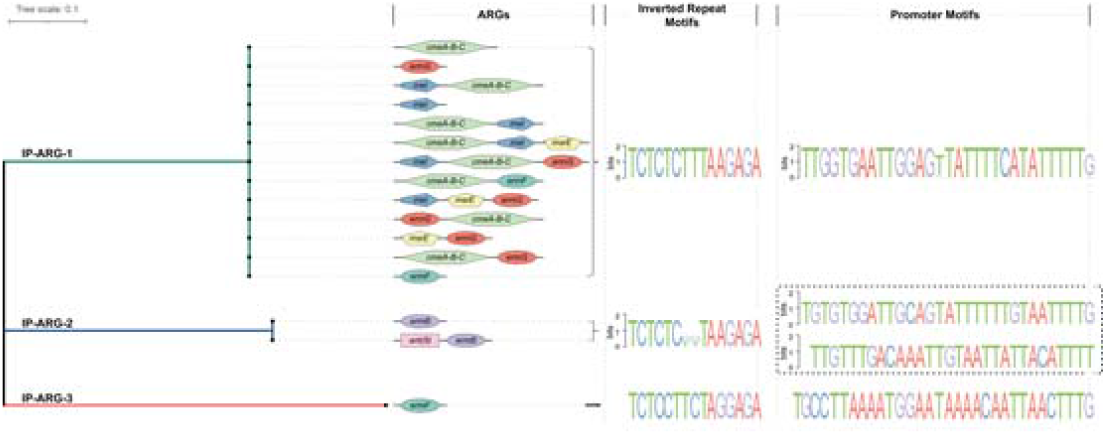
Phase-variable ARGs were classified into three classes (IP-ARG-1, IP-ARG-2, and IP-ARG-3) based on the alignment of nucleotide sequences of invertase genes. The ARGs and putative ARG organization patterns are shown for each class. Different ARGs are shown in different shapes and colors.

To further characterize these three classes of phase-variable ARGs, we analyzed the motifs of inverted repeats and invertible promoters as well as the ARGs regulated by the invertible regions. The invertible promoter motifs varied among classes but were nearly identical within the same class (Figure 1). The inverted repeat motifs were found to be similar across different classes (Figure 1). Interestingly, we found two invertible regions located immediately downstream of the invertase gene in class IP-ARG-2 (Figure 1). The two invertible regions were located adjacent to each other and the inverted repeats were nearly identical in these two regions. This suggested that these two invertible regions could be regulated by the same upstream invertase gene. However, the invertible promoters were different from each other (Figure 1, Table S2). This suggested that these two invertible regions might generate more phenotypic variations than only one invertible region does. The ARGs regulated by invertible promoters varied among classes (Figure 1, Table S2). Most invertible promoters of class IP-ARG-1 regulated the *cmeABC* operon, *ermG* gene, or both, while some also regulated other ARGs, including *mel, msrE, tetQ*, or *ermF*. These ARGs could confer resistance to diverse antibiotics, including cephalosporin, fluoroquinolone, fusidic acid, lincosamide, macrolides, oxazolidinone, phenicol, pleuromutilin, streptogramin, and tetracycline. The *ermB* gene and the *ermF* gene, conferring resistance to streptogramin, macrolides, and lincosamides, were found to be regulated by class IP-ARG-2 and class IP-ARG-3, respectively. In addition, class IP-ARG-2 was also found to regulate an *ant(9)* gene homolog which confers resistance to aminoglycosides.

### Taxonomic distribution of phase-variable ARGs

To examine the geographic and taxonomic distribution of phase-variable ARGs, we searched genomes from the Unified Human Gastrointestinal Genome (UHGG) collection (29) to detect nearly identical (> 99% identity) invertase genes. The invertase genes mediating ARGs were grouped into the corresponding class of phase-variable ARGs (Table S3). Phase-variable ARGs were observed in multiple countries, but the geographic prevalence varied across classes (Table 1, Table S2, and Table S3). Phase-variable ARGs from classes IP-ARG-1 and IP-ARG-2 were identified in metagenomes from 17 and 7 countries, respectively, spanning three continents (Asia, Europe, and North America). Phase-variable ARGs from class IP-ARG-3 were only observed in the metagenomes from Denmark, which might be due to the limited sampling of publically-available metagenomic data. The results reveal that species harboring phase-variable ARGs are widely geographically distributed.

**Table 1.** Taxonomic and geographic distribution of the phase-variable ARGs.

| The class of identified<br>Integrase genes and<br>identical homologs<br>regulating ARGs | Bacteria host <sup>1</sup> |  |  |  | Country |
| --- | --- | --- | --- | --- | --- |
|  | Order | Family | Genus | Species |  |
| 1 | Bacteroidales | Bacteroidaceae | Bacteroides | Bacteroides cellulosilyticus, Bacteroides eggerthii, Bacteroides fluxus, Bacteroides fragilis, Bacteroides fragilis_A, Bacteroides intestinalis, Bacteroides ovatus, Bacteroides stercoris, Bacteroides thetaiotaomicron, Bacteroides uniformis, Bacteroides xylanisolvens, Unclassified species | Austria, Canada, China, Denmark, Estonia, Finland, France, Germany, Israel, Japan, Kazakhstan, Netherlands, Russia, Spain, Sweden, United Kingdom, United States |
|  |  |  | Phocaeicola | Phocaeicola coprocola, Phocaeicola coprophilus, Phocaeicola dorei, Phocaeicola sp000432735, Phocaeicola vulgatus |  |
|  |  |  | Prevotella | Prevotella copri, Prevotella lascolaii, Prevotella sp000834015, Prevotella sp001275135, Prevotella sp900313215, Prevotella stercorea |  |
|  |  | Barnesiellaceae | Barnesiella | Barnesiella intestinihominis |  |
|  |  | Coprobacteraceae | Coprobacter | Coprobacter fastidiosus |  |
|  |  | Tannerellaceae | Parabacteroides | Parabacteroides distasonis, Parabacteroides goldsteinii, Parabacteroides merdae |  |
| 2 | Bacteroidales | Bacteroidaceae | Bacteroides | Bacteroides caccae, Bacteroides eggerthii, Bacteroides fragilis, Bacteroides thetaiotaomicron, Bacteroides uniformis | China, Denmark, Estonia, Israel, Italy, United Kingdom, United States |
|  |  |  | Paraprevotella | Paraprevotella xylaniphila |  |
|  |  |  | Phocaeicola | Phocaeicola dorei |  |
| 3 | Bacteroidales | Bacteroidaceae | Phocaeicola | Phocaeicola dorei | Denmark |
|  |  | UBA932 | RC9 | RC9 sp000434935 |  |

Phase-variable ARGs were found to be commonly and exclusively distributed in *Bacteroidales* species (Table 1, Table S2, and Table S3). Phase-variable ARGs belonging to class IP-ARG-1 were observed in 28 *Bacteroidales* species from the families *Bacteroidaceae, Barnesiellaceae, Coprobacteraceae*, and *Tannerellaceae*. Phase-variable ARGs belonging to class IP-ARG-2 could be identified in 6 species from *Bacteroidaceae* that belonged to the order *Bacteroidales*. Class IP-ARG-3 was found in two species, *Phocaeicola dorei* and a novel species RC9 sp000434935, which also belongs to the *Bacteroidales* order. These suggested that phase-variable ARGs are restricted in *Bacteroidales*. The wide taxonomic distribution combined with the sparse occurrence of these phase-variable ARGs is not consistent with vertical transmission, suggesting that phase-variable ARGs could be horizontally transferred by mobile genetic elements (MGEs).

### All three classes of phase-variable ARGs are located within integrative conjugative elements

To determine whether the phase-variable ARGs were on MGEs, we searched the invertases and invertible regions regulating phase-variable ARGs as well as the flanking sequences against ImmeDB (30) and the ICEberg database (31). We found that all three classes of phase-variable ARGs are located within integrative conjugative elements (ICEs) (FIgure 2). The majority of phase-variable ARGs from class IP-ARG-1 were identified within ICEs related to ICE26 in the ImmeDB (Figure 2a). Phase-variable ARGs from class IP-ARG-1 were also found within another novel ICE (Figure 2a). This suggested that the progenitor of IP-ARG-1 might have emerged before subsequently inserting into multiple ICEs. Phase-variable ARGs from class IP-ARG-2 were detected in ICEs related to ICE14 in ImmeDB (Figure 2b). A phase-variable ARG from class IP-ARG-3 contained within a ∼50-kb sequence fragment most closely matches ICE34 in ImmeDB (Figure 2c). The fact that all three classes of phase-variable ARGs are located within ICEs is medically relevant because ICEs encode the necessary machinery to horizontally transfer between species mobilizing any ARGs that they acquire.

**Figure 2:**
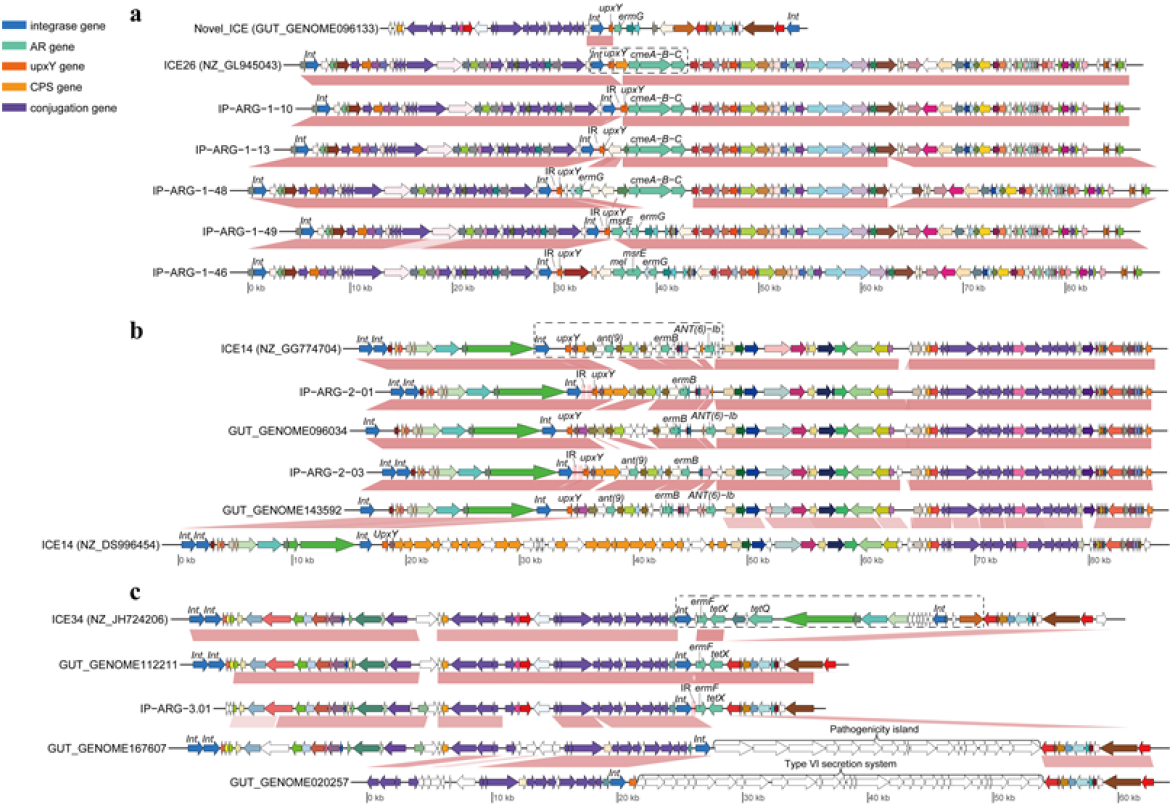
Genomic comparison and context analysis of the integrative and conjugative element (ICE) variants carrying IP-ARG-1 (a), IP-ARG-2 (b), or IP-ARG-3 (c). The regions located adjacent to the invertible promoters (black dotted boxes) were found to be highly variable across element variants within each ICE. ICE26, ICE14, and ICE34 are ICE accession numbers from ImmeDB. The NCBI genome accession numbers are shown in the parentheses after the ICE accession numbers. The sequence labels that start with GUT_GENOME are genome accession numbers of the UHGG database. Orthologous genes are plotted with the same color and are linked by pink connections. Invertase and antibiotic resistance genes are colored blue and light green, respectively. Genes involved in conjugation and capsular polysaccharide biosynthesis are colored purple and orange, respectively. The genes that do not have orthologs are white. Invertase, *upxY*, and antibiotic resistance genes as well as inverted repeats are labeled. AR, CPS, INV, and IR denote antibiotic resistance, capsular polysaccharides, invertase, and inverted repeats, respectively.

Phase-variable ARGs were frequently observed to be included in a highly variable region that was located immediately downstream of the invertible regions. The highly varied regions contained genes that were not necessary for the ICE replication and transfer, but often important for conferring selective and adaptive advantages for hosts in the changing environments (Figure 2). Most of the variable genes in the highly varied region were ARGs in ICEs containing class IP-ARG-1 (Figure 2a, Table S2), which suggested multiple insertions of ARGs at this region. In ICEs carrying class IP-ARG-2, not only the ARGs but also the CPS genes, even clusters that contained only CPS genes, were located in the regions downstream of the invertases (Figure 2b). In different ICEs with class IP-ARG-3, the highly variable regions contained varied genes or operons, such as ARGs, operons involved in pathogenicity island or T6SS, or integrative and mobilizable elements encoding ARG *tetQ* (Figure 2c). All these indicated that the loci in such highly variable regions downstream of the invertible promoters were hotspots for the acquisition of elements, including ARGs.

### The invertible regions regulating ARGs appear to originate from those regulating CPS genes independently and convergently

To better understand the evolution of phase-variable ARGs, we performed comparative analyses on the highly varied regions regulated by invertible promoters. We found that CPS genes, such as *wecA* gene, or the genes necessary for CPS operon, like *upxY* gene (32), were frequently located immediately downstream of the invertible promoters and upstream of the identified ARGs (Figure 2a, b, Figure 3). This suggested that the invertible promoters regulating ARGs and those regulating CPS related genes may share the same origin. In addition, the inverted repeats and the invertase amino acids in class IP-ARG-2 shared 100% and more than 96% identity with the corresponding sequences of those regulating CPS clusters, respectively (Figure 2b). These results support that the invertible regions regulating ARGs share a common evolutionary origin with invertible regions regulating CPS genes.

**Figure 3:**
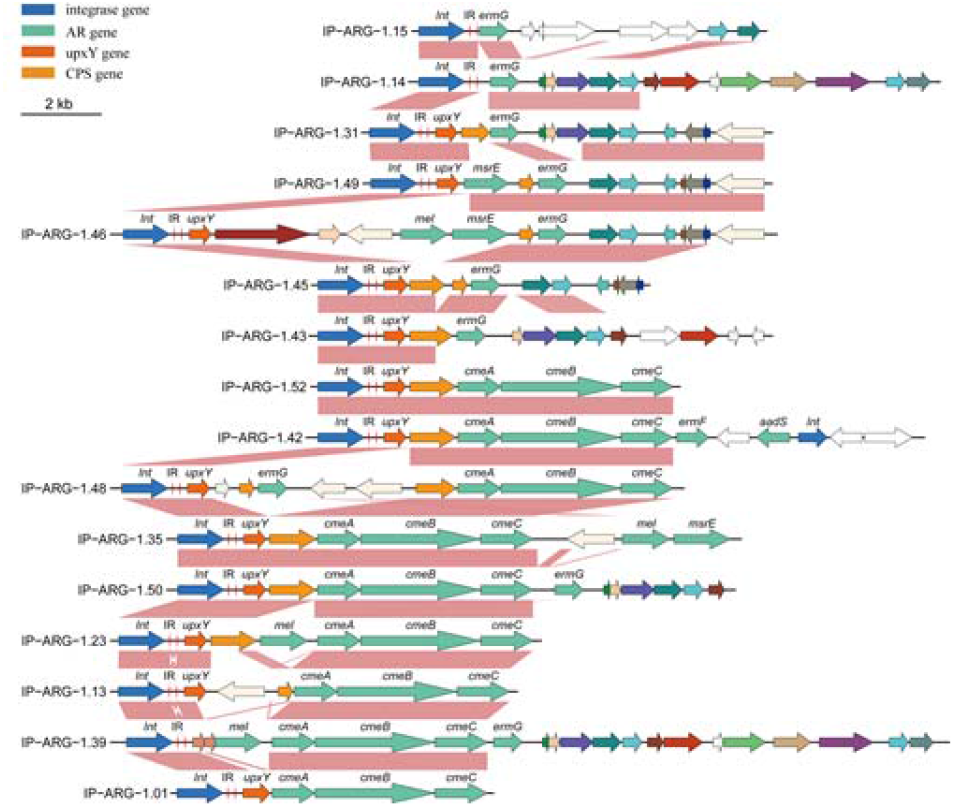
Comparisons of highly variable regions carrying IP-ARG-1 demonstrate the degeneration of the *upxY* gene. Different degeneration statuses of the *upxY* gene were: partially degenerated (IP-ARG-1-39), completely degenerated (IP-ARG-1-14 and IP-ARG-1-15), fusion gene (IP-ARG-1-01), and intact gene (the remainder). Orthologous genes are plotted with the same color and are linked by pink connections. Invertase genes and antibiotic resistance genes are colored blue and light green, respectively. Genes involved in conjugation and capsular polysaccharide biosynthesis are colored purple and orange, respectively. The genes that do not have orthologs are white. Invertase, *upxY*, and antibiotic resistance genes as well as inverted repeats are labeled. AR, CPS, INV, and IR denote antibiotic resistance, capsular polysaccharides, invertase, and inverted repeats, respectively.

**Figure 4:**
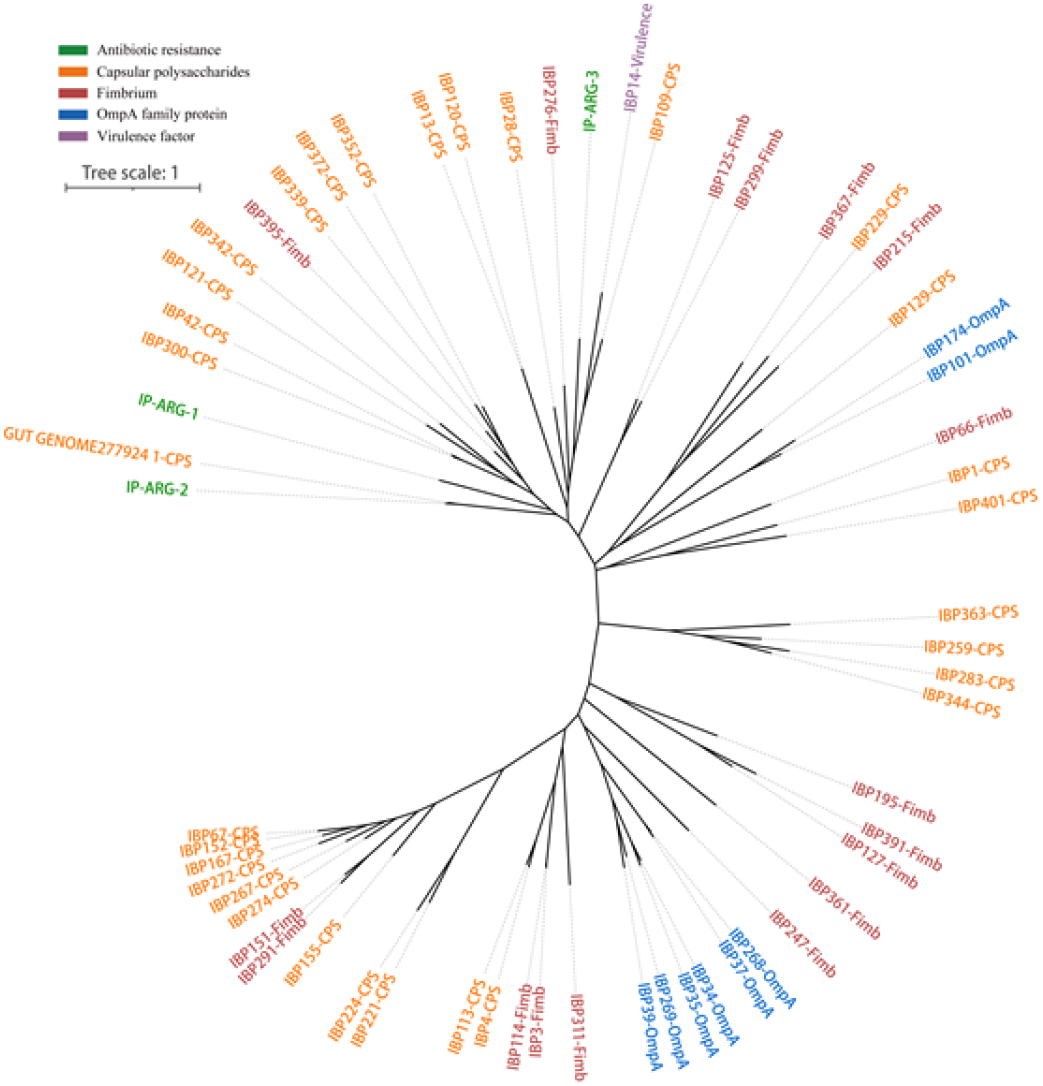
The evolutionary events that led to the emergence of phase-variable ARGs may have occurred independently at least three times as the result of convergent evolution. The phylogenetic tree was inferred based on the alignment of nucleotide sequences of invertase genes. The invertase genes include those mediating phase-variable ARGs identified in our assembled sequences and their homologs in the UHGG database as well as those that have been previously reported(17). IBP denotes the invertible DNA region in phylum Bacteroidetes, which is designated in the previous study(17). The labels that start with GUT_GENOME are genome accessions in the UHGG database. Labels are colored based on the functional annotation of the loci regulated by the invertase genes and invertible regions.

Gene context analysis further shows that invertible regions regulating ARGs appeared to be derived from invertible regions originally regulating CPS genes. The phase-variable ARGs were frequently found to be located both upstream and downstream of CPS genes (Figure 2a, b). The genes downstream of the invertible promoters were often partially degenerated, completely degenerated, or fused with the partial sequence of the *wecA* gene (Figure 2a).

To understand the evolutionary history of different classes of IP-ARGs, a phylogenetic tree was constructed based on the nucleotide sequences of the local invertase genes of known invertible promoters in Bacteroidetes (Figure 3). The result suggests that the evolutionary events that led to the emergence of different classes of phase-variable ARGs occurred independently. Class IP-ARG-1 might have emerged before being mobilized by ICEs. Class IP-ARG-2 might have emerged as the ARG inserted into the CPS gene cluster loci that had been carried by ICEs (Figure 2a, b). It was difficult to infer the evolutionary history of class IP-ARG-3 because we have identified few sequences closely related to this class. However, the phylogenetic tree revealed that IP-ARG-3 and another two classes were distributed in distinct clades, suggesting that the emergence of class IP-ARG-3 was likely independent of class IP-ARG-1 or IP-ARG-2. Hence, the emergence of different classes of phase-variable ARGs might be the result of convergent evolution and the evolutionary events that led to the emergence of phase-variable ARGs likely occurred independently at least three times.

## DISCUSSION

In this study, we systematically searched for phase-variable ARGs in human gut microbes and explored the origin, evolution, and prevalence of the phase-variable ARGs. Our analysis revealed that phase-variable ARGs are (a) commonly found in *Bacteroidales* species, (b) often originate from ARG insertions into phase-variable CPS loci, (c) frequently mobilized by ICEs which may explain their wide taxonomic distribution within the phylum Bacteroidetes and their rapid dissemination, and (d) widely geographically distributed.

Our analysis only identified phase-variable ARGs in *Bacteroidales* species. There could be several reasons for this observed taxonomic restriction. First, gut *Bacteroidales* genomes typically contain numerous phase-variable loci. In some *Bacteroides* species, such as *Bacteroides fragilis*, there are more than 20 phase-variable loci including up to 7 phase-variable CPS loci (33, 34). While other phyla prevalent in the gut, including Proteobacteria, Firmicutes, and Actinobacteria, have phase-variable loci, these loci are far less common and have far fewer examples per genome (17, 35). Therefore, ARG insertions into Bacteroidetes phase-variable loci are statistically more likely than insertions into phase-variable loci in other phyla. Furthermore, as abundant members of the human gut microbiome, metagenomic sequencing leads to high coverage of Bacteroidetes species which increases the likelihood that a phase-variable region can be detected with PhaseFinder (17). Though more than two thousand metagenomic samples were analyzed, no phase-variable ARGs were identified in other phyla, suggesting phase-variable ARGs might be rare in phyla other than Bacteroidetes.

This finding highlights the threat to human health that Bacteroidetes species pose as a reservoir for the dissemination of ARGs. The phylum Bacteroidetes is one of the most abundant in the human gut and Bacteroidetes species, especially those in genus *Bacteroides*, are regarded as a reservoir of antibiotic resistance genes (36). Moreover, members of Bacteroidetes species, such as *Bacteroides fragilis*, are considered opportunistic pathogens (37) and can be the causative agent of appendicitis and intra-abdominal abscesses (38, 39). Antibiotics have been used to treat such infections, but an increasing rate of antibiotic resistance has been noted in Bacteroidetes species (40–43). Phase-variable ARGs may contribute to the continued maintenance of clinically-relevant antibiotic resistance genes such as *cmeABC, ermG*, and *tetQ*, which confer resistance to a widely-used antibiotics including macrolides, streptogramin and tetracycline (42, 44). Most species identified with phase-variable ARGs are considered commensals in the human gut. However, ARGs transfer from commensals to pathogens through horizontal transfer has been documented (5, 45, 46). The transfer of phase-variable ARGs from commensals to pathogens via MGEs, such as the ICEs identified in this study, might promote resistance to a wide array of antibiotics in pathogenic species (47), posing a threat to public health in the future. Due to the fact that inter-phylum transfer of ICEs is rare (30), the spread of phase-variable ARGs might be limited within the phylum Bacteroidetes. As such, monitoring these Bacteroidetes species, especially the ICEs in these species, might be important in mitigating the threat of phase-variable ARGs to human health.

## MATERIALS AND METHODS

### De novo assembly and gene annotation of metagenomic datasets

We downloaded metagenomic sequencing data that consisted of 2227 human gut samples encompassing 7 studies (Table S1). Low-quality reads were removed and sequencing adapters were trimmed with trim_galore (v0.6.4; https://github.com/FelixKrueger/TrimGalore). The filtered data were mapped to the human genome (hg19) using bowtie2 (v2.3.5.1) (48) to filter human reads. The cleaned reads from each sample were assembled with SPAdes (v3.14.0) using the - -meta option (49, 50), and sequences less than 500 bp were removed. Gene annotation was performed using Prokka (v1.14.5) (51) with the parameter --metagenome.

### Annotation of antibiotic resistance genes (ARGs)

We searched the annotated sequences for known ARGs from the Comprehensive Antibiotic Resistance Database (CARD) (v3.0.7) (52) using Blastn (v2.10.0) (53). The BLAST results were filtered using the parameters: -perc_identity 80, -evalue 1e-10, and -culling_limit 1. Resistance Gene Identifier (RGI, v5.0.0) (52) was also used to predict known antibiotic resistance elements using the following parameters: rgi main --t contig -a BLAST -n 8 -d wgs --local. In addition, we also searched the sequences against the Resfams HMM Database (Core) (54) using hmmscan with the parameter: --cut_ga.

### Identification of phase-variable ARGs

ARGs, along with upstream and downstream 20,000 flanking bases, were extracted to identify invertons using PhaseFinder (v1.0) (17). The default parameters of PhaseFinder were used. We filtered the results by removing the invertible DNA regions with < 2 reads supporting the R orientation from the paired-end method, and the Pe_ratio < 1%. Furthermore, the invertible DNA regions containing or overlapping coding sequences (CDS) were removed. The motifs of inverted repeats and promoters were identified based on the previously reported motifs (17) and the logos were generated with WebLogo (version 2.8.2) (55).

### Identification of host species and mobile genetic elements

We identified the host species for the identified contigs using blastn by searching the non-MGE sequences against the NCBI non-redundant nucleotide (nt) database and the Unified Human Gastrointestinal Genome UHGG database (29) with an e-value <1-e10. We performed blastn to search for identical homologs (> 99% identity) of the identified invertases and invertible regions against the 204,938 non-redundant genomes from the UHGG database. The identical homologs were grouped into corresponding classes of IP-ARG and the UHGG annotations for host species were examined to identify host species for IP-ARG.

To identify integrative and conjugative elements (ICEs) in different classes of IP-ARGs, we searched the ICEberg (31, 56) and ImmeDB (30) databases using blastn with an e-value <1-e10. We used ConjScan (57, 58) via a Galaxy web server (https://galaxy.pasteur.fr) to annotate conjugative genes in ICEs.

### Genomic comparison and phylogenetic analysis

The bacteria genomes and genome comparisons were visualized with the R package genoPlotR. All identified invertases were used to construct the phylogenetic tree for the classification of IP-ARGs. To understand the evolutionary history of IP-ARGs, the invertase genes regulating phase-variable ARGs identified in our assembled sequences and their homologs in the UHGG database, as well as those that have been previously reported(17) were used to construct the phylogenetic tree. Redundant invertase genes were filtered using cd-hit with the 99% identity threshold (59). Only the invertase genes regulating functionally characterized genes or operons were included in the tree. Multiple alignments of invertase gene nucleotide sequences were performed with MUSCLE (v3.8.31) (60). The alignment results were analyzed in FastTree (v2.1.10) (61) with default parameters to infer the phylogenetic trees. The phylogenetic trees were visualized using iTOL (https://itol.embl.de).

## Acknowledgments

We thank NIH’s Biowulf cluster team. This work utilized the computational resources of the NIH HPC Biowulf cluster (http://hpc.nih.gov).

W.Y. and X.J. is supported by the Intramural Research Program of the NIH, National Library of Medicine. B.H. is supported by startup funding from the University of Maryland.

## Contributions

X.J. conceived and designed the study. W.Y. performed the data analysis and wrote the draft manuscript. X.J. and A.B.H. reviewed and revised the manuscript. X.J. supervised the work. All authors read and approved the final manuscript.

## Competing interests

The authors declare no competing interests.

## Supplementary Information

### Supplementary Data

Supplementary Data 1: Nucleotide sequences of the identified contigs carrying phase-variable ARGs.

### Supplementary Tables

Table S1: Human gut metagenomic samples analyzed in this study.

Table S2: Sequences containing phase-variable antibiotic resistance genes.

Table S3: UHGG genomes carrying the homologs of the identified invertase genes mediating phase-variable ARGs.

## Notes

### Competing Interest Statement

The authors have declared no competing interest.

